# RulePep: Interpretable ESM-Guided Neural-Symbolic Peptide Classification

**DOI:** 10.64898/2026.07.03.736448

**Authors:** Farzad Midjani, Reza Ghelich, Fateme Zahra Keshtkar, Mahdi Malekpour, Heewook Lee

## Abstract

Peptides are increasingly explored as therapeutic candidates, delivery vectors, and functional biomolecules, but experimental screening of peptide activity and safety remains costly because the sequence space is vast and small sequence changes can alter functionality. Computational peptide classification can therefore help prioritize candidates. However, many protein-language-model-based classifiers achieve strong performance using opaque prediction heads, making it difficult to determine which learned evidence supports or opposes a prediction. We present RulePep, an ESM-2-guided neural-symbolic classifier for peptide-function prediction. RulePep maps frozen ESM-2 sequence representation to learned latent predicates, polarity-constrained differentiable rules, and an additive symbolic logit whose components can be inspected at the case level. We evaluate RulePep on three biologically distinct peptide classification tasks: blood-brain barrier penetration, hemolytic potency, and anticancer activity. On the BBPpredict, HemoPI3, and AntiCP 2.0 alternate benchmark datasets, RulePep achieved AUROC/MCC values of 0.8869/0.6850, 0.9155/0.6820, and 0.9765/0.8633, respectively. Ablation experiments supported the contributions of multi-layer representation pooling, rule polarity, mined-rule initialization, symbolic capacity, and rule-derived aggregation. RulePep combines competitive predictive performance with additive logit reconstruction, rule-level evidence reporting, and predicate-suppression auditing, providing a transparent sequence-based framework for peptide candidate prioritization.

## 1 Introduction

Peptides are an important class of biological molecules with growing relevance in therapeutic discovery, molecular delivery, and functional screening. Their relatively small size, sequence programmability, and ability to mediate selective molecular or membrane interactions make them attractive candidates for drug development and biotechnology. At the same time, peptide activity is highly sequence dependent: changes in change distribution, hydrophobicity, aromatic content, amphipathicity, or local motifs can substantially alter biological function, toxicity, and stability. Because the possible peptide sequence space is enormous, experimental testing alone cannot exhaustively evaluate candidate peptides. Sequence-based machine learning can therefore serve as an important triage step by prioritizing candidates before synthesis and experimental evaluation.

This need arises across several biologically distinct peptide-design problems. lood–brain barrier (BBB) penetration is a major obstacle for central nervous system drug delivery, and BBB-penetrating peptides may help identify molecules or carrier sequences capable of reaching neural tissue. Hemolytic potency is an important safety endpoint because membrane-active peptides can damage red blood cells, limiting their therapeutic utility even when they show desirable antimicrobial or anticancer activity. Anticancer peptides are of interest as potential therapeutic agents because some peptides can selectively disrupt cancer cells or interfere with tumor-associated biological processes. Although BBB penetration, hemolysis, and anticancer activity represent different biological properties, each can be formulated as a binary peptide-classification task in which the input is an amino-acid sequence and the output is a task-specific activity label. This shared formulation allows a common sequence-modeling architecture to be trained and evaluated across functionally different peptide benchmarks.

Transfer learning and large-scale sequence pretraining have established protein language models (PLMs) as effective biological sequence encoders [1, 2]. Models such as ProteinBERT [3], ProtTrans [4], ESM [5], and peptide-specific transformer models [6] have extended this paradigm to protein and peptide property prediction. However, many peptide classifiers still attach dense, recurrent, convolutional or ensemble prediction heads to PLM embeddings. Such models may perform well, but their final scores are often difficult to decompose into positive and negative evidence. This limitation is important in peptide discovery, where users may want to understand not only whether a candidate is predicted to be active, but also which learned sequence-derived evidence supports activity, which evidence raises safety concerns, and how competing signals affect the final score.

Neural-symbolic learning can provide more transparent sequence classifiers by introducing explicit rule-like structures into trainable models, including neural-symbolic reasoning foundations [7], differentiable logic layers [9, 11], probabilistic logic programming [10], and rule learning from noisy data [8]. We therefore investigate whether a shared PLM-based neural-symbolic architecture can support multiple peptide-classification endpoints while retaining a decomposable decision path.

We present RulePep, a neural-symbolic classifier that maps frozen ESM-2 representations to learned latent predicates, expands those predicates into presence and absence literals, initializes polarity-constrained rule banks, and combines direct and higher-order rule-derived evidence into an additive symbolic logit. We evaluate the proposed model on three benchmark peptide-classification tasks: BBPpredict benchmark dataset [12], the HemoPI3 benchmark dataset [13], and the AntiCP 2.0 alternate benchmark dataset [25].

## 2 Materials and Methods

Figure 1 summarizes the three-stage pipeline: ESM-based representation learning, symbolic rule induction, and prediction with evidence reporting.

**Figure 1.**
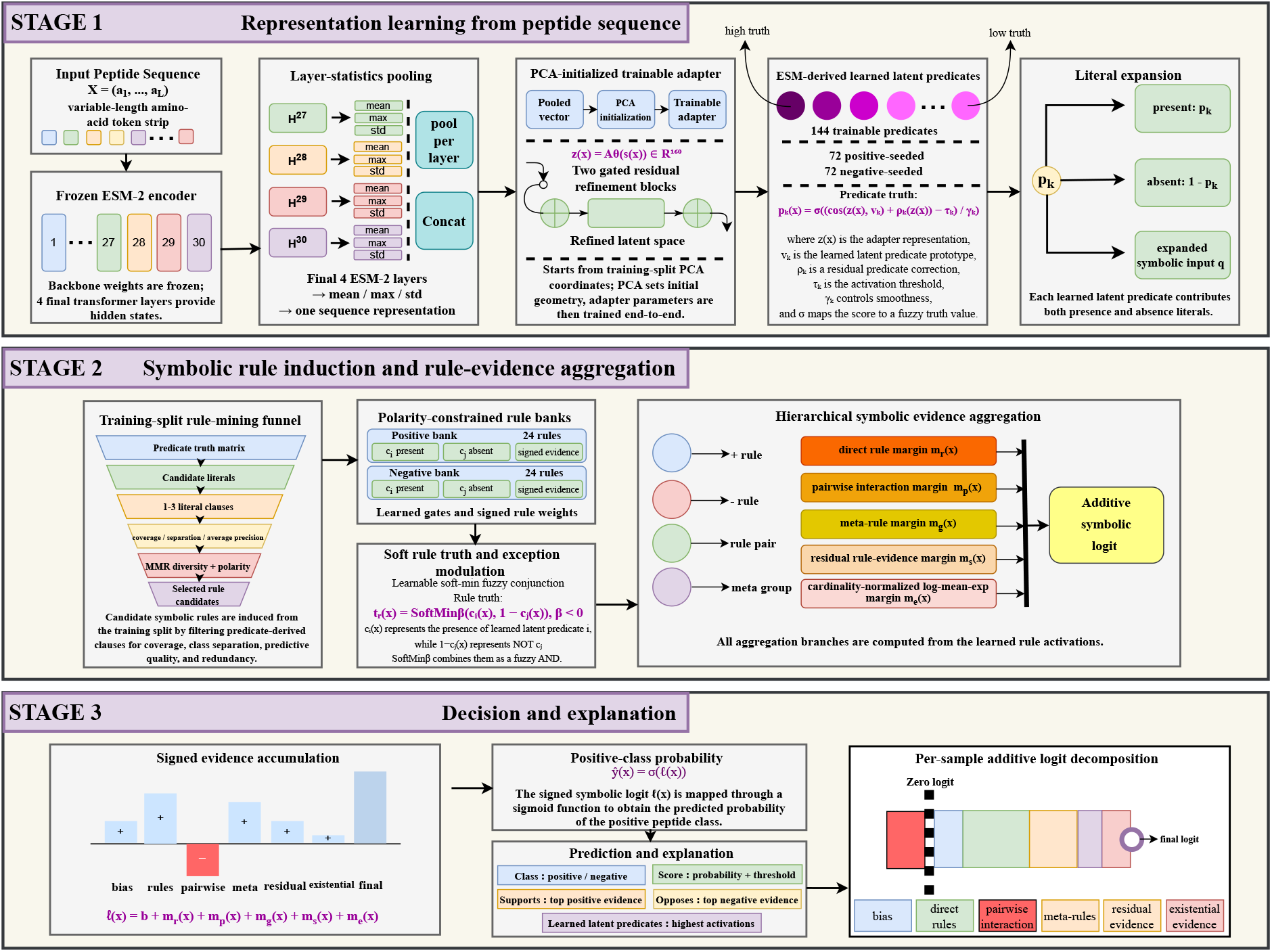
Architecture of RulePep. Final-layer ESM-2 representations are pooled, adapted, and mapped to 144 learned latent predicates. Training-split mining initializes polarity-specific rule banks whose direct and higher-order terms form the final logit. The displayed two-literal equation is a simplified illustration of the implemented one-to three-literal rules. Rule-bank sizes denote initialization capacity before validation-based pruning.

### 2.1 Problem definition and data handling

All three case studies are treated as binary peptide-classification tasks. For a peptide sequence 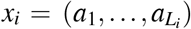, the binary label *y*_*i*_ ∈ {0, 1} specifies class membership for the corresponding task: *y*_*i*_ = 1 represents the task-specific positive class, namely BBB-penetrating peptides in BBPpredict, highly hemolytic peptides in HemoPI3, and anticancer peptides in the AntiCP 2.0 alternate dataset, while *y*_*i*_ = 0 represents the corresponding negative class. The same model architecture, representation pipeline, symbolic classifier, and evaluation procedure are used across the three datasets; the training sequences and label semantics differ by task.

Preprocessing was limited to task-agnostic sequence cleaning. Non-standard residues were mapped to X, and sequences outside the 2-200 residue range were removed. Exact duplicates were collapsed, and conflicting duplicate sequences were excluded. The final cleaned dataset sizes were 844 sequences for BBPpredict (421 positive, 423 negative), 1623 sequences for HemoPI3 (885 positive, 738 negative), and 1940 sequences for the AntiCP 2.0 alternate dataset (970 positive, 970 negative).

Across the three case studies, the cleaned data are partitioned once using a 70/15/15 train/validation/test design with class-stratified splitting. The training partition fits the ESM adapter, learned latent predicates, differentiable rules, rule interactions, rule-derived aggregation branches, and train-split rule-mining initialization. The validation partition is reserved for checkpoint, threshold, and pruning decisions, and the held-out test partition is evaluated after these choices have been fixed.

### 2.2 ESM representation, adapter, and learned latent predicates

A frozen ESM-2 encoder, `esm2_t30_150M_UR50D`, maps each peptide into contextual residue representations [5]. Let *H*^(*k*)^ ∈ ℝ^*L×d*^ denote the ESM representation at transformer layer *k*. The representation uses the final four layers, *K* = {27, 28, 29, 30}, and forms a sequence vector by concatenating residue-wise mean, maximum, and standard-deviation statistics over these layers,

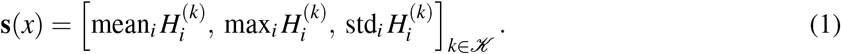

The predicate layer maps the ESM-2 representation to trainable fuzzy activation values. These latent predicates are initialized from class-conditional representation structure and do not correspond to human-annotated biological concepts. The pooled training representations are standardized using training-split statistics and projected by principal component analysis (PCA) into a 160-dimensional initialization space [20]. Let 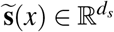 denote the standardized form of the pooled representation in Eq. 1. The trainable adapter produces the refined latent representation

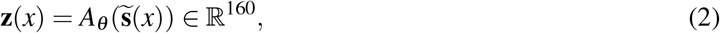

where *A*_*θ*_ denotes the PCA-initialized adapter with parameters *θ*. At initialization,

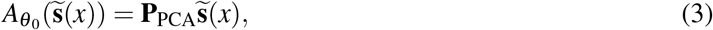

where 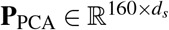 was fitted only to the training representations. The adapter contains two gated residual blocks with zero-initialized output layers; it therefore begins as the PCA projection and subsequently learns nonlinear corrections jointly with the symbolic classifier.

RulePep contains 144 learned latent predicates arranged as 72 positive-seeded and 72 negative-seeded predicates. In each bank, 47 prototypes were initialized by training-split *k*-means centroids and 25 by diverse class-boundary examples [21]. Boundary examples were ranked by proximity to the opposite-class centroid, distance from their own-class centroid, and redundancy. Predicate *k* has prototype *v*_*k*_, threshold *τ*_*k*_, temperature *γ*_*k*_, and residual correction *ρ*_*k*_:

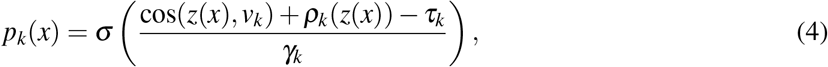

where *ρ*_*k*_(*z*) is a trainable residual predicate score regularized toward zero. All predicate and adapter parameters are then optimized jointly with the symbolic classifier; the rule layer therefore receives trainable fuzzy predicate values rather than fixed cluster assignments.

### 2.3 Predicate literals and train-split rule induction

The learned latent predicate vector is expanded into a literal vector containing both predicate-presence and predicate-absence literals,

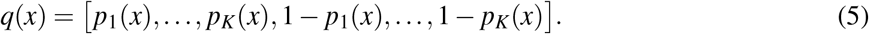

The model maintains separate positive and negative rule banks. Positive rules may select positive-seeded predicate-presence and negative-seeded predicate-absence literals; negative rules may select negative-seeded predicate-presence and positive-seeded predicate-absence literals. This constraint preserves the class-oriented initialization and ensures that positive-bank and negative-bank rules contribute with predefined logit directions.

Candidate clauses containing one to three literals were mined exclusively from the training split. Eligible clauses required class separation ≥0.05, average precision ≥0.60, target-class coverage ≥0.05, and at least 15 activated training samples. Maximum marginal relevance with *λ*_MMR_ = 0.45 selected nonredundant clauses using rule-truth similarity and literal overlap [22]. The selected clauses initialized 24 positive and 24 negative rules, whose gates, confidences, literal-selection logits, and non-negative evidence weights were optimized end-to-end.

Rule *r* is evaluated using the learnable negative-order generalized mean in Eq. 6. Here, *a*_*r j*_ = *σ*(*α*_*r j*_) denotes the selection strength of literal *j*, and the learned *β* < 0 produces a differentiable soft-min aggregation dominated by the least-satisfied selected literal, consistent with fuzzy conjunction [23]:

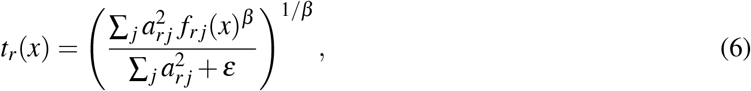

where

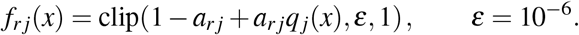

The exponent is parameterized as

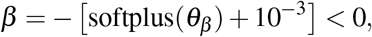

yielding a differentiable approximation to the minimum of the selected literal values. Squaring *a*_*r j*_ reduces the influence of nearly inactive literals. Rules without an effectively selected literal are assigned the neutral conjunction value, and the final rule truth is clipped to [*ε*, 1 −*ε*].

### 2.4 Symbolic scoring, conflict handling, and rule-derived aggregation branches

Let 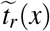 denote the rule truth after cross-polarity exception modulation. For positive-class-supporting rules *ℛ*^*+*^ and negative-class-supporting rules *ℛ*^−^, the direct rule margin is

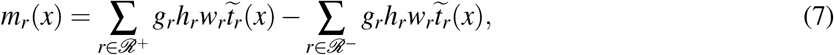

where *g*_*r*_ is the rule gate, *h*_*r*_ is the rule confidence, and *w*_*r*_ ≥ 0 is the contribution scale. The bank assignment determines the sign of each contribution; the learned weights themselves are non-negative.

In addition to the direct rule margin, the classifier contains four rule-derived components: signed pairwise interactions, positive- and negative-class meta-rule groups, a residual aggregation branch, and a cardinality-normalized, temperature-scaled log-mean-exp aggregation of class-specific active-rule supports. All four operate on the same exception-modulated rule activations rather than on an independent neural prediction head.

For direction *d* ∈ {+, −}, let 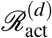 denote the rules that remain active after validation-based pruning and optional intervention masking. The confidence-weighted support of rule *r* is

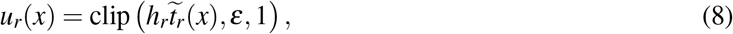

where *h*_*r*_ is the learned rule confidence and 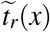 is the exception-modulated rule truth. The normalized class-specific log-support is

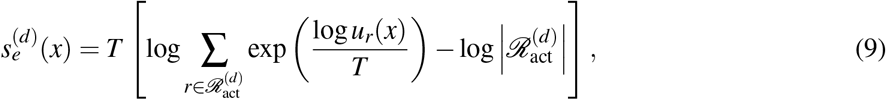

where *T* > 0 is the existential aggregation temperature. The existential margin is

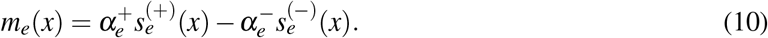

The normalization removes the additive rule-count term produced by unnormalized log-sum-exp aggregation. Because each normalized class-specific log-support is non-positive, the combined margin *m*_*e*_(*x*), rather than either term alone, determines whether the existential branch increases or decreases the positive-class logit.

The final logit is the additive sum of the symbolic components used during inference:

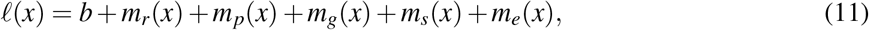

where *m*_*p*_ denotes the pairwise rule-interaction margin, *m*_*g*_ the gated meta-rule margin, *m*_*s*_ the residual rule-evidence aggregation margin, and *m*_*e*_ the cardinality-normalized existential log-support contrast defined in Eq. 10. The positive-class probability is 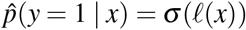.

### 2.5 Training objective and model selection

The training objective combines class-weighted binary cross-entropy with ranking, evidence-alignment, branch-supervision, predicate-regularization, rule-regularization, sparsity, and logit-stability terms:

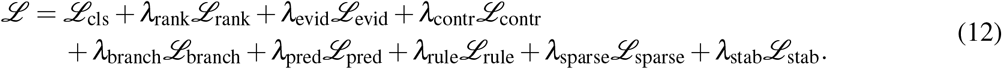

Here, *ℒ*_rank_ is a pairwise logit-ranking term; *ℒ*_evid_ aligns positive and negative evidence with the corresponding labels; *ℒ*_contr_ penalizes simultaneous positive and negative evidence; and *ℒ*_branch_ supervises the residual and existential rule-aggregation margins. *ℒ*_pred_ combines predicate-group separation, contradiction, diversity, residual-correction, and usage penalties. *ℒ*_rule_ combines rule-diversity, rule-coverage, polarity-balance, and conjunction-regularization terms. *ℒ*_sparse_ regularizes selection, confidence, exception, interaction, meta-rule, and binarization parameters, whereas *ℒ*_stab_ controls logit magnitude. During training, rule activations were subjected to dropout with probability 0.015 [24]. Models were implemented in PyTorch [18] and optimized with AdamW. Experiments used a maximum sequence length of 200 residues, ESM batch size of 8, validation-selected checkpointing, and gradient clipping. Post-training pruning was accepted only when the validation composite score decreased by no more than 0.002. When pruning was accepted, the retained bank contained at most 22 rules, including at least four positive and four negative rules when available. If pruning failed this criterion, the unpruned 48-rule model was retained. Accepted pruned models were fine-tuned for at most 20 epochs with patience 8 and a learning-rate multiplier of 0.35; otherwise, the pre-pruning model was restored.

### 2.6 Evaluation and comparison protocol

Performance was evaluated using accuracy, precision, recall (sensitivity), specificity, F1-score, AUROC, and Matthews correlation coefficient (MCC).

### 2.7 Ablation configurations

Ten ablation variants were evaluated on the BBPpredict dataset using the same data split, optimization, validation-based checkpoint selection, and threshold-selection procedure as the full model. Representation ablations used only the final ESM layer or halved each predicate bank from 72 to 36 predicates. Rule-capacity ablations halved each rule bank from 24 to 12 rules, disabled rule dropout, removed polarity constraints, removed residual or existential aggregation, replaced the symbolic classifier with a neural head, disabled mined-rule initialization, or used product conjunction without exceptions, pairwise interactions, or meta-rules.

### 2.8 Evidence reporting and decision-path auditing

For each peptide, the model reports the predicted positive-class probability, the additive components of Eq. 11, the activated predicates and rules, and the changes produced by individual rule removal.

Decision-path auditing comprised algebraic reconstruction and rule removal. Algebraic reconstruction compared the inference logit with

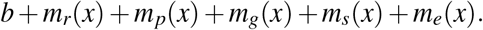

For rule *r*, the total removal effect was

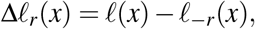

where all rule-dependent branches were recomputed after disabling *r*. Its dependency-mediated component was

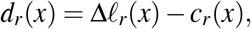

with *c*_*r*_(*x*) denoting the rule’s direct additive contribution.

### 2.9 Sequence-level predicate suppression and residue-window localization

For each test peptide, the ordinary pooled ESM-2 representation was passed through the trained adapter and predicate layer to obtain the sequence-level predicate vector **p**(*x*) = [*p*_1_(*x*), …, *p*_*K*_(*x*)]. The predicates selected for intervention were the eight predicates with the highest sequence-level truth values for that peptide.

For an intervention on predicate *j*, its sequence-level truth value was suppressed according to

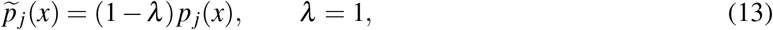

while all other predicate values and all trained parameters were held fixed. The modified predicate vector was then passed through the same literal-expansion, rule-activation, evidence-aggregation, logit, and probability computations used during standard inference. The signed intervention effect was defined as

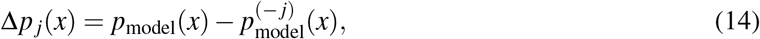

where 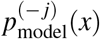 is the probability after suppressing predicate *j*.

Residue-level scores were computed separately for localization. For residue *i*, a proxy representation was constructed from the final four ESM-2 layers:

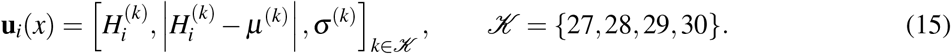

After standardization with the training-split scaler, **u**_*i*_(*x*) was passed through the trained adapter and predicate function to obtain a token-level proxy truth *p* _*j,i*_(*x*). Presence literals were localized by maximizing *p* _*j,i*_(*x*), whereas absence literals were localized by maximizing 1 − *p* _*j,i*_(*x*). A radius-two residue window was displayed around the maximizing position:

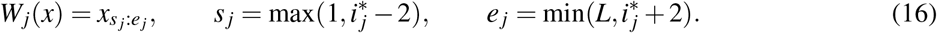

Token-level proxy scores were used to identify residue windows; all baseline and post-suppression probabilities were computed from sequence-level predicate values. The windows therefore represent proxy associations with predicate activation.

## 3 Results and Discussion

### 3.1 Held-out performance across peptide functions

RulePep achieved held-out MCC values of 0.6850, 0.6820, and 0.8633 on the BBPpredict, HemoPI3, and AntiCP 2.0 alternate datasets, respectively (Table 1).

**Table 1.** Held-out test performance of RulePep across the three benchmark datasets.

| Benchmark dataset | Accuracy | AUROC | Precision | Recall | F1 | MCC |
| --- | --- | --- | --- | --- | --- | --- |
| BBPpredict dataset | 0.8425 | 0.8869 | 0.8413 | 0.8413 | 0.8413 | 0.6850 |
| HemoPI3 dataset | 0.8410 | 0.9155 | 0.8713 | 0.8302 | 0.8502 | 0.6820 |
| AntiCP 2.0 alternate dataset | 0.9313 | 0.9765 | 0.9496 | 0.9103 | 0.9296 | 0.8633 |

*Note: Values are reported as the means of three independent runs with different random seeds. The standard deviation across runs did not exceed 0.02 for any reported metric.*

### 3.2 Learned symbolic rule-bank composition

Validation retained all 48 initialized rules for the BBPpredict, HemoPI3 and AntiCP 2.0 alternate datasets. Figure 2 details the BBPpredict rule bank, and Table 2 summarizes rule-bank composition across all three benchmark datasets.

**Table 2.** Cross-dataset composition of mined candidates and learned symbolic rule banks.

| Benchmark dataset | Mined candidates | Selected initial rules | Final active rules | Positive rule bank | Negative rule bank |
| --- | --- | --- | --- | --- | --- |
| BBPpredict dataset | 500 (16/151/333) | 48 (6/29/13) | 48 (6/29/13) | 24 (4/12/8; 34/18) | 24 (2/17/5; 14/37) |
| HemoPI3 dataset | 500 (34/408/58) | 48 (20/14/14) | 48 (20/14/14) | 24 (9/11/4; 20/23) | 24 (11/3/10; 27/20) |
| AntiCP 2.0 alternate dataset | 500 (23/310/167) | 48 (13/30/5) | 48 (13/30/5) | 24 (5/18/1; 25/19) | 24 (8/12/4; 17/27) |

*Note:* Parenthetical triplets denote the numbers of one-, two-, and three-literal clauses, respectively. In the positive- and negative-rule-bank columns, the pair after the semicolon denotes the numbers of presence and absence literals, respectively.

**Figure 2.**
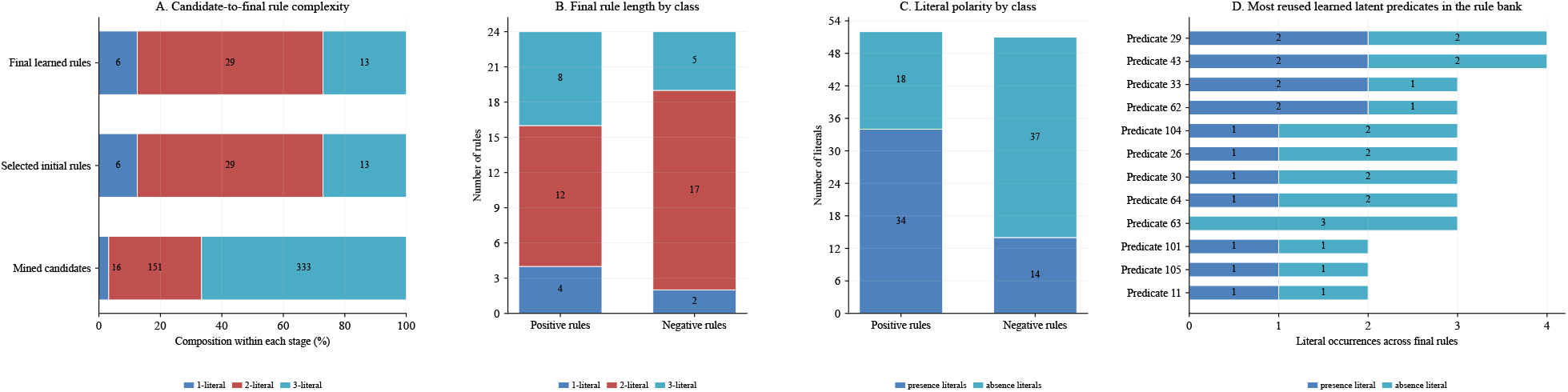
Symbolic rule-bank composition on BBPpredict. (A) Clause lengths among the top 500 eligible candidates and 48 validation-retained rules. (B) Rule lengths by positive and negative banks. (C) Presence and absence literals by bank. (D) Most frequently reused learned latent predicates. Labels show raw counts.

### 3.3 Additive-logit reconstruction and contribution-intervention concordance

Additive-logit reconstruction and rule-removal agreement were evaluated for the final validation-selected model on each benchmark dataset (Table 3).

**Table 3.**
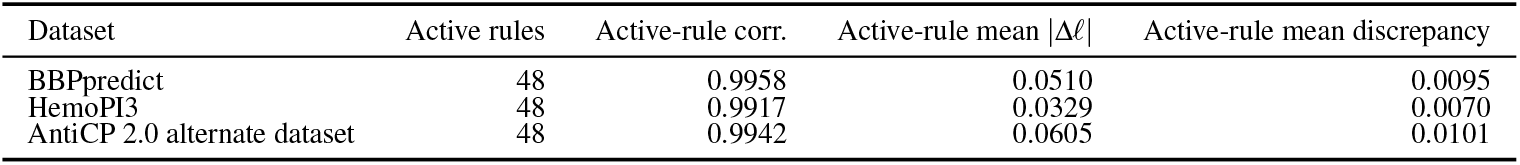
Decision-path checks for the final validation-selected neural-symbolic classifier.

| Dataset | Active rules | Active-rule corr. | Active-rule mean $\vert \Delta\ell\vert $ | Active-rule mean discrepancy |
| --- | --- | --- | --- | --- |
| BBPpredict | 48 | 0.9958 | 0.0510 | 0.0095 |
| HemoPI3 | 48 | 0.9917 | 0.0329 | 0.0070 |
| AntiCP 2.0 alternate dataset | 48 | 0.9942 | 0.0605 | 0.0101 |

Figures 3 and 4 show, respectively, case-level logit decomposition and predicate-suppression localization for representative BBPpredict test samples.

**Figure 3.**
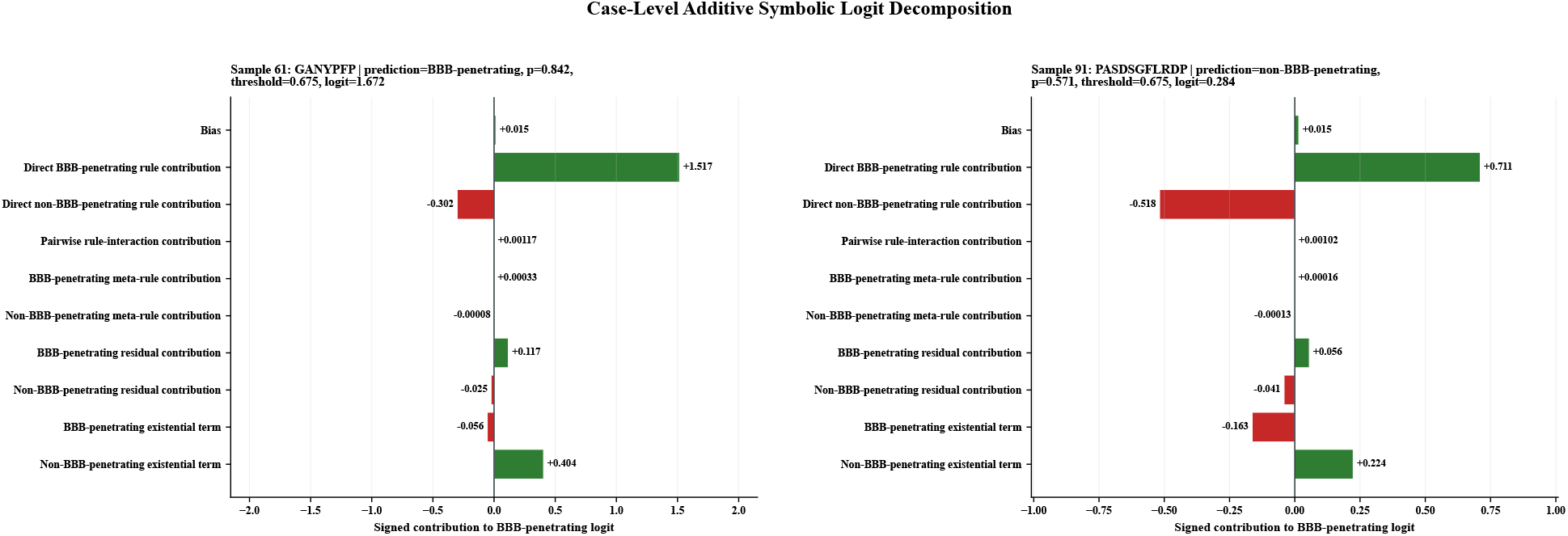
Case-level decomposition of the BBB-penetrating logit for two BBPpredict test peptides. Bars show signed contributions from all additive components. Sample 61 was classified as BBB-penetrating 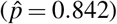, whereas Sample 91 was classified as non-BBB-penetrating because 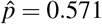 was below the validation-selected threshold of 0.675. Contributions sum to the reported logits up to rounding.

**Figure 4.**
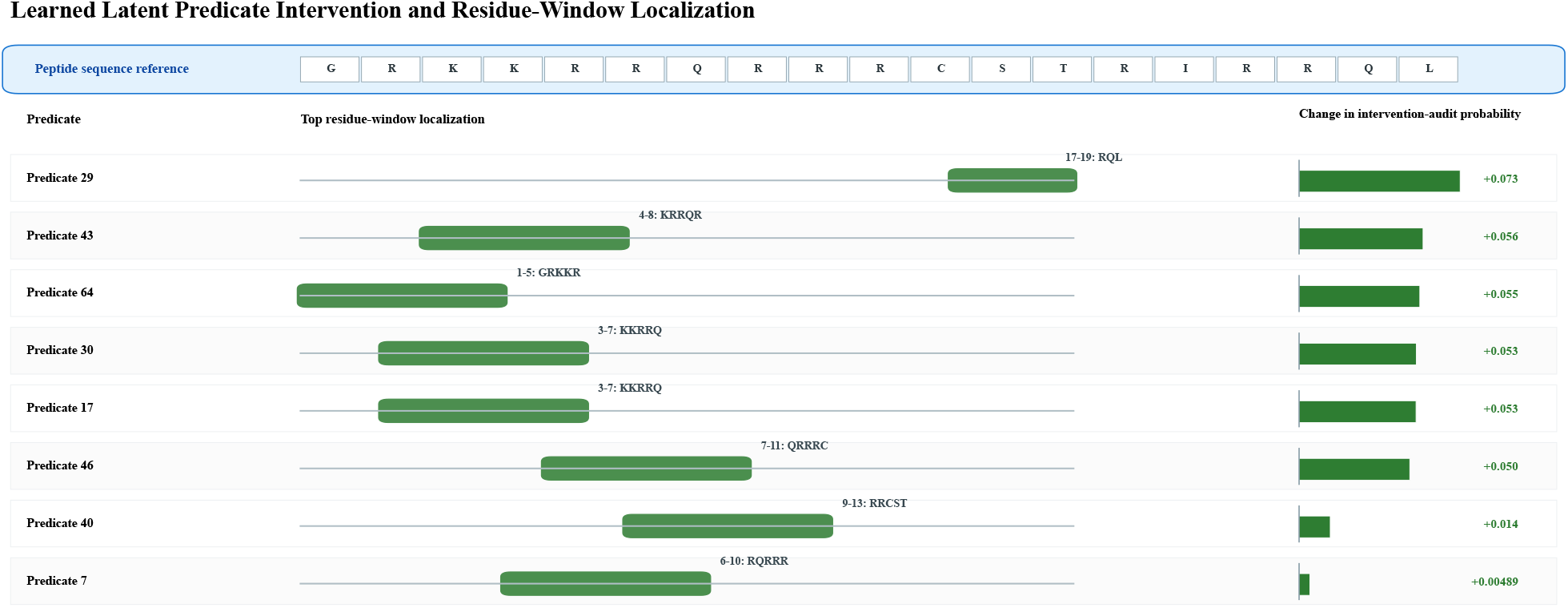
Predicate-suppression audit with residue-window localization for BBPpredict test peptide (sample 67). For each predicate, Δ*p* = *p*_baseline_ *p*_suppressed_ was computed after setting its sequence-level truth to zero. Residue windows indicate the highest-scoring radius-two token-level localization.

Figure 5 summarizes sequence-level predicate-suppression effects for the proposed model on the BBPpredict benchmark dataset. Across the three benchmark datasets, the intervention analysis evaluated only predicates that appeared among the eight highest-truth predicates for at least one test peptide.

**Figure 5.**
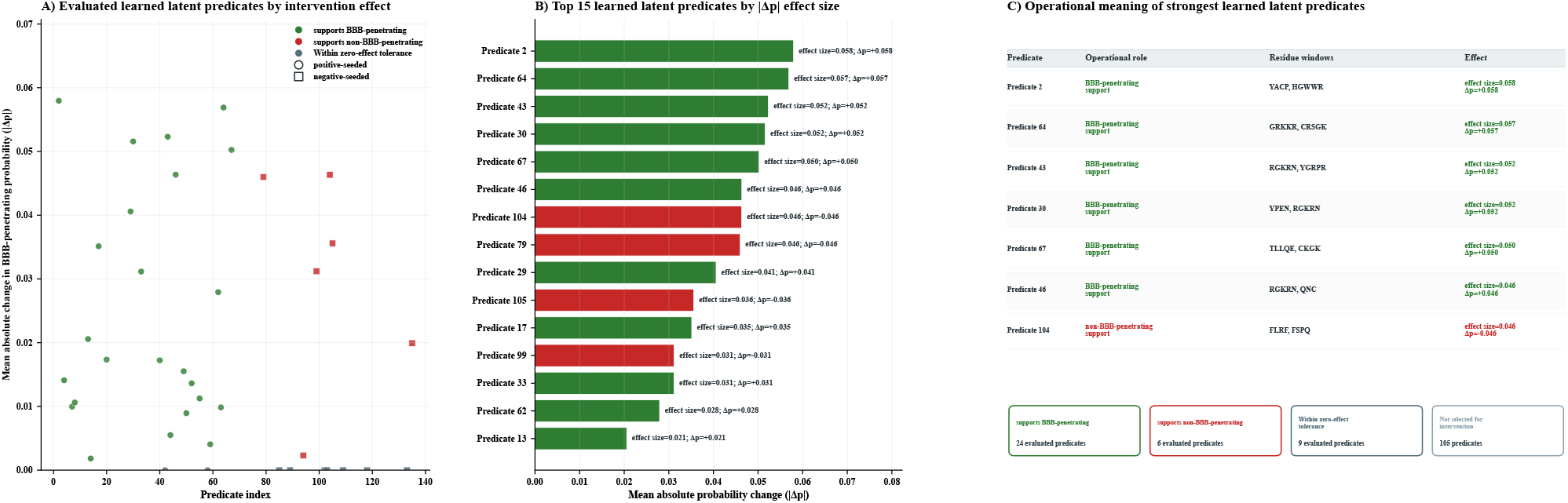
Predicate-suppression analysis for the proposed model on the BBPpredict dataset. (A) Mean absolute probability changes across learned latent predicates, with colors indicating signed effects and marker shapes indicating initialization. (B) The 15 predicates with the largest Δ*p*, where Δ*p* = −*p*_baseline_ *p*_suppressed_. (C) Operational roles, representative residue windows, and effects of the strongest predicates. Of 144 predicates, 39 were evaluated.

### 3.4 Ablation analysis

Table 4 compares the full model with ten matched ablations on the BBPpredict benchmark dataset.

**Table 4.** Matched ablation results on the BBPpredict independent test partition. Values are means across the same three training seeds used to evaluate the full model.

| Configuration | Accuracy | AUROC | Precision | Recall | MCC | ΔMCC |
| --- | --- | --- | --- | --- | --- | --- |
| Full model | 0.8425 | 0.8869 | 0.8413 | 0.8413 | 0.6850 | 0.0000 |
| Final ESM layer only | 0.7647 | 0.8447 | 0.7872 | 0.7255 | 0.5310 | -0.1540 |
| Compact learned latent predicate bank | 0.7745 | 0.8677 | 0.7917 | 0.7451 | 0.5500 | -0.1350 |
| Shallow rule base | 0.7647 | 0.8285 | 0.7872 | 0.7255 | 0.5310 | -0.1540 |
| No symbolic rule dropout | 0.8137 | 0.8812 | 0.8077 | 0.8235 | 0.6276 | -0.0574 |
| No strict rule polarity | 0.7549 | 0.8562 | 0.7407 | 0.7843 | 0.5107 | -0.1743 |
| No residual evidence-aggregation branch | 0.7451 | 0.8085 | 0.7451 | 0.7451 | 0.4902 | -0.1948 |
| No existential evidence-aggregation branch | 0.7549 | 0.8401 | 0.7708 | 0.7255 | 0.5107 | -0.1743 |
| Neural-only head | 0.7549 | 0.8589 | 0.7826 | 0.7059 | 0.5123 | -0.1727 |
| No mined-rule initialization | 0.7953 | 0.8795 | 0.7846 | 0.8095 | 0.5909 | -0.0941 |
| Minimal symbolic layer | 0.7638 | 0.8752 | 0.7538 | 0.7778 | 0.5279 | -0.1571 |

All ten ablations had lower MCC than the full model on the fixed BBPpredict test split. The largest reductions followed removal of the residual aggregation branch (ΔMCC = −0.1948), strict rule polarity (−0.1743), the existential aggregation branch (−0.1743), and the symbolic classifier (−0.1727). These point estimates identify polarity constraints and rule-derived aggregation as the most influential tested components under this split.

### 3.5 Contextual comparison with published benchmark results

Table 5 compares the proposed model with source-reported benchmark results for the corresponding peptide-classification tasks.

**Table 5.** Benchmark comparator metrics for the three peptide-classification case studies. Each comparator row is attributed to the source paper that reported or introduced the corresponding model.

| Benchmark case study | Comparator model | Source | Acc. | Rec. | Spec. | MCC |
| --- | --- | --- | --- | --- | --- | --- |
| *BBPPredict: BBB-penetrating vs. non-BBB peptide prediction* |  |  |  |  |  |  |
| BBPPredict dataset | RulePep | This work | 0.8425 | 0.8413 | 0.8438 | 0.6850 |
| BBPPredict dataset | BBB-PEP-prediction | [16] | 0.8240 | 0.8310 | 0.9110 | 0.6630 |
| BBPPredict dataset | BBPPredict RF | [12] | 0.7727 | 0.7677 | 0.7778 | 0.5455 |
| BBPPredict dataset | BBPPredict rbfSVM | [12] | 0.7626 | 0.7879 | 0.7374 | 0.5259 |
| BBPPredict dataset | LogitBoost | [12] | 0.7273 | 0.6768 | 0.7778 | 0.4569 |
| BBPPredict dataset | GentleBoost | [12] | 0.7071 | 0.7475 | 0.6667 | 0.4155 |
| *HemoPI3: highly hemolytic vs. poorly hemolytic peptide prediction* |  |  |  |  |  |  |
| HemoPI3 dataset | RulePep | This work | 0.8410 | 0.8302 | 0.8539 | 0.6820 |
| HemoPI3 dataset | HemoPI dipeptide-composition SVM | [13] | 0.8238 | 0.8193 | 0.8306 | 0.6400 |
| HemoPI3 dataset | HEPAD-XGBoost | [17] | 0.8060 | 0.8250 | 0.7840 | 0.6090 |
| HemoPI3 dataset | CNN sequence model | [19] | 0.8031 | 0.8294 | - | 0.6051 |
| HemoPI3 dataset | HemoPred | [15, 17] | 0.7980 | 0.8500 | 0.7330 | 0.5900 |
| HemoPI3 dataset | HLPpred-Fuse | [14] | 0.7920 | 0.8210 | 0.7620 | 0.5850 |
| *AntiCP 2.0 alternate: anticancer vs. non-anticancer peptide prediction* |  |  |  |  |  |  |
| AntiCP 2.0 alternate dataset | RulePep | This work | 0.9313 | 0.9103 | 0.9521 | 0.8633 |
| AntiCP 2.0 alternate dataset | AntiC P2.0 AAC-ETree | [25] | 0.9201 | 0.9227 | 0.9175 | 0.8400 |
| AntiCP 2.0 alternate dataset | ACP-MHCNN | [28, 30] | 0.9000 | 0.8650 | 0.9330 | 0.8000 |
| AntiCP 2.0 alternate dataset | AI4ACP | [26] | 0.8940 | 0.8710 | 0.9180 | 0.7900 |
| AntiCP 2.0 alternate dataset | iACP-FSCM | [26, 27] | 0.8890 | 0.8760 | 0.9020 | 0.7790 |
| AntiCP 2.0 alternate dataset | ACP-DL | [28, 29] | 0.8300 | 0.8290 | 0.8290 | 0.6600 |

Note: Comparator values are source-reported results obtained under study-specific evaluation protocols. Some results originate from later studies that re-evaluated publicly available implementations; in these cases, both the original model paper and the re-evaluation source are cited.

The RulePep point estimates compare favorably with several historical comparator values in Table 5, while differences in data partitioning, threshold selection, and evaluation protocols should be considered.

### 3.6 Implications and limitations

The reported contributions are computed directly from the inference path and describe how the model forms its score. They do not establish biochemical mechanisms. The current evidence is also limited by random sequence-level splitting, incomplete calibration analysis, validation-selected thresholds, and the absence of external experimental validation.

## 4 Conclusion

We proposed an ESM-guided neural-symbolic classifier for binary peptide-function prediction that combines frozen PLM representations, learned latent predicates, differentiable rules, and logit-level evidence reconstruction. Our results further show that frozen PLM embeddings can be decomposed into a set of interpretable latent predicates that support symbolic decision rules without compromising predictive performance.

## 5 Code and Data Availability

The source code supporting the findings of this study is publicly available in the RulePep GitHub repository at https://github.com/Farzad-Midjani/RulePep, and the benchmark datasets used in the three case studies are publicly available through the resources associated with the original studies.

## References

[1] Rao R, Bhattacharya N, Thomas N, Duan Y, Chen X, Canny J, Abbeel P, Song YS. Evaluating protein transfer learning with TAPE. In: Advances in Neural Information Processing Systems. 2019;32:9689–9701.

[2] Rives A, Meier J, Sercu T, Goyal S, Lin Z, Liu J, et al. Biological structure and function emerge from scaling unsupervised learning to 250 million protein sequences. Proceedings of the National Academy of Sciences. 2021;118(15):e2016239118. doi:10.1073/pnas.2016239118.

[3] Brandes N, Ofer D, Peleg Y, Rappoport N, Linial M. ProteinBERT: a universal deep-learning model of protein sequence and function. Bioinformatics. 2022;38(8):2102–2110. doi:10.1093/bioinformatics/btac020.

[4] Elnaggar A, Heinzinger M, Dallago C, Rehawi G, Wang Y, Jones L, et al. ProtTrans: Toward Understanding the Language of Life Through Self-Supervised Learning. IEEE Transactions on Pattern Analysis and Machine Intelligence. 2022;44(10):7112–7127. doi:10.1109/TPAMI.2021.3095381.

[5] Lin Z, Akin H, Rao R, Hie B, Zhu Z, Lu W, et al. Evolutionary-scale prediction of atomic-level protein structure with a language model. Science. 2023;379(6637):1123–1130. doi:10.1126/science.ade2574.

[6] Guntuboina C, Das A, Mollaei P, Kim S, Barati Farimani A. PeptideBERT: a language model based on transformers for peptide property prediction. The Journal of Physical Chemistry Letters. 2023;14(46):10427–10434. doi:10.1021/acs.jpclett.3c02398.

[7] d’Avila Garcez AS, Lamb LC, Gabbay DM. Neural-Symbolic Cognitive Reasoning. Berlin, Heidelberg: Springer; 2009. doi:10.1007/978-3-540-73246-4.

[8] Evans R, Grefenstette E. Learning explanatory rules from noisy data. Journal of Artificial Intelligence Research. 2018;61:1–64. doi:10.1613/jair.5714.

[9] Badreddine S, d’Avila Garcez A, Serafini L, Spranger M. Logic Tensor Networks. Artificial Intelligence. 2022;303:103649. doi:10.1016/j.artint.2021.103649.

[10] Manhaeve R, Dumančić S, Kimmig A, Demeester T, De Raedt L. DeepProbLog: Neural probabilistic logic programming. In: Advances in Neural Information Processing Systems. 2018;31:3749–3759.

[11] Serafini L, d’Avila Garcez AS. Logic Tensor Networks: Deep Learning and Logical Reasoning from Data and Knowledge. arXiv preprint. 2016;arXiv:1606.04422.

[12] Chen X, Zhang Q, Li B, Lu C, Yang S, Long J, et al. BBPpredict: A web service for identifying blood-brain barrier penetrating peptides. Frontiers in Genetics. 2022;13:845747. doi:10.3389/fgene.2022.845747.

[13] Chaudhary K, Kumar R, Singh S, Tuknait A, Gautam A, Mathur D, Anand P, Varshney GC, Raghava GPS. A web server and mobile app for computing hemolytic potency of peptides. Scientific Reports. 2016;6:22843. doi:10.1038/srep22843.

[14] Hasan MM, Schaduangrat N, Basith S, Lee G, Shoombuatong W, Manavalan B. HLPpred-Fuse: improved and robust prediction of hemolytic peptide and its activity by fusing multiple feature representation. Bioinformatics. 2020;36(11):3350–3356. doi:10.1093/bioinformatics/btaa160.

[15] Win TS, Malik AA, Prachayasittikul V, Wikberg JES, Nantasenamat C, Shoombuatong W. HemoPred: A web server for predicting the hemolytic activity of peptides. Future Medicinal Chemistry. 2017;9(3):275–291. doi:10.4155/fmc-2016-0188.

[16] Naseem A, Alturise F, Alkhalifah T, Khan YD. BBB-PEP-prediction: improved computational model for identification of blood-brain barrier peptides using blending position relative composition specific features and ensemble modeling. Journal of Cheminformatics. 2023;15(1):110. doi:10.1186/s13321-023-00773-1.

[17] Chen SH, Yu JC, Lin YH, Kuo SC, Ni K, Chen CT. HEPAD: enhancing hemolytic peptide prediction with adaptive feature engineering and diverse sequence descriptors. BMC Bioinformatics. 2025;26:234. doi:10.1186/s12859-025-06254-6.

[18] Paszke A, Gross S, Massa F, Lerer A, Bradbury J, Chanan G, et al. PyTorch: An imperative style, high-performance deep learning library. In: Advances in Neural Information Processing Systems. 2019;32:8024–8035.

[19] Abdelbaky I, Elhakeem M, Tayara H, Badr E, Abdul Salam M. Enhanced prediction of hemolytic activity in antimicrobial peptides using deep learning-based sequence analysis. BMC Bioinformatics. 2024;25(1):368. doi:10.1186/s12859-024-05983-4.

[20] Jolliffe IT, Cadima J. Principal component analysis: a review and recent developments. Philosophical Transactions of the Royal Society A. 2016;374(2065):20150202. doi:10.1098/rsta.2015.0202.

[21] MacQueen J. Some methods for classification and analysis of multivariate observations. In: Proceedings of the Fifth Berkeley Symposium on Mathematical Statistics and Probability. Vol. 1. Berkeley: University of California Press; 1967. p. 281–297.

[22] Carbonell J, Goldstein J. The use of MMR, diversity-based reranking for reordering documents and producing summaries. In: Proceedings of the 21st Annual International ACM SIGIR Conference on Research and Development in Information Retrieval. 1998. p. 335–336. doi:10.1145/290941.291025.

[23] Klement EP, Mesiar R, Pap E. Triangular Norms. Dordrecht: Springer; 2000. doi:10.1007/978-94-015-9540-7.

[24] Srivastava N, Hinton G, Krizhevsky A, Sutskever I, Salakhutdinov R. Dropout: a simple way to prevent neural networks from overfitting. Journal of Machine Learning Research. 2014;15:1929–1958.

[25] Agrawal P, Bhagat D, Mahalwal M, Sharma N, Raghava GPS. AntiCP 2.0: an updated model for predicting anticancer peptides. Briefings in Bioinformatics. 2021;22(3):bbaa153. doi:10.1093/bib/bbaa153.

[26] Sun YY, Lin TT, Cheng WC, Lu IH, Lin CY, Chen SH. Peptide-based drug predictions for cancer therapy using deep learning. Pharmaceuticals. 2022;15(4):422. doi:10.3390/ph15040422.

[27] Charoenkwan P, Chiangjong W, Lee VS, Nantasenamat C, Hasan MM, Shoombuatong W. Improved prediction and characterization of anticancer activities of peptides using a novel flexible scoring card method. Scientific Reports. 2021;11:3017. doi:10.1038/s41598-021-82513-9.

[28] Ahmed S, Muhammod R, Khan ZH, Adilina S, Sharma A, Shatabda S, Dehzangi A. ACP-MHCNN: an accurate multi-headed deep-convolutional neural network to predict anticancer peptides. Scientific Reports. 2021;11:23676. doi:10.1038/s41598-021-02703-3.

[29] Yi HC, You ZH, Zhou X, Cheng L, Li X, Jiang TH, Chen ZH. ACP-DL: a deep learning long short-term memory model to predict anticancer peptides using high-efficiency feature representation. Molecular Therapy-Nucleic Acids. 2019;17:1–9. doi:10.1016/j.omtn.2019.04.025.

[30] Liu M, Wu T, Li X, Zhu Y, Chen S, Huang J, Zhou F, Liu H. ACPPfel: explainable deep ensemble learning for anticancer peptides prediction based on feature optimization. Frontiers in Genetics. 2024;15:1352504. doi:10.3389/fgene.2024.1352504.

